# Intrinsic functional architecture of the macaque dorsal and ventral lateral frontal cortex

**DOI:** 10.1101/058776

**Authors:** Alexandros Goulas, Peter Stiers, R. Matthew Hutchison, Stefan Everling, Michael Petrides, Daniel S. Margulies

**Affiliations:** Max Planck Research Group for Neuroanatomy and Connectivity, Max Planck Institute for Human Cognitive and Brain Sciences, Leipzig, Germany; Faculty of Psychology and Neuroscience, Department of Neuropsychology and Psychopharmacology, Maastricht University, Maastricht, The Netherlands; Center for Brain Sciences, Harvard University, Cambridge, MA, USA; Robarts Research Institute, University of Western Ontario, London, Ontario, Canada; Cognitive Neuroscience Unit, Montreal Neurological Institute, McGill University, Montreal, Quebec, Canada

**Keywords:** cortical areas, in-vivo parcellation, functional connectivity, premotor, prefrontal

## Abstract

Investigations of the cellular and connectional organization of the lateral frontal cortex (LFC) of the macaque monkey provide indispensable knowledge for generating hypotheses about the human LFC. However, despite numerous investigations, there are still debates on the organization of this brain region. In vivo neuroimaging techniques such as resting-state fMRI can be used to define the functional circuitry of brain areas producing results largely consistent with gold-standard invasive tract-tracing techniques and offering the opportunity for cross-species comparison within the same modality. Our results using resting-state fMRI from macaque monkeys to uncover the intrinsic functional architecture of the LFC corroborate previous findings and inform current debates. Specifically, we show that i) the region in the midline and anterior to the superior arcuate sulcus is divided in two areas separated by the posterior supraprincipal dimple; ii) the cytoarchitectonically defined area 6DC/F2 contains two connectional divisions; and, iii) a distinct area occupies the cortex around the spur of the arcuate, updating what was previously proposed to be the border between dorsal and ventral motor/premotor areas. Within the ventral LFC specifically, the derived parcellation clearly suggests the presence of distinct areas i) with a somatomotor/orofacial connectional signature (putative area 44), ii) with an occulomotor connectional signature (putative frontal eye fields), and iii) premotor areas possibly hosting laryngeal and arm representations. Our results illustrate in detail the intrinsic functional architecture of the macaque LFC, thus providing valuable evidence for debates on its organization.

## Introduction

Cytoarchitectonic and myeloarchitectonic investigations of the macaque monkey lateral frontal cortex (LFC) have provided critical information on its organization (e.g., Vogt and Vogt, 1919; Walker, 1940; Barbas and Pandya, 1987; Petrides and Pandya, 1994). Furthermore, investigations of the cortico-cortical connections of these areas with invasive tract-tracing methods have provided evidence of distinct connectivity profiles that characterize these cytoarchitectonically distinct areas (e.g., Cavada and Goldman-Rakic, 1989; Petrides and Pandya, 2006; Yeterian et al.,2012). Thus, cytoarchitectonic and connectional investigations have unveiled a mosaic of cortical areas within the LFC (Figure 1) that participate in specific large-scale networks. In addition, electrophysiological recordings in these areas and selective lesion studies have provided evidence of relative functional specializations of the neuronal populations in these cortical areas (e.g. Petrides, 2005; Kaping et al., 2011). Despite considerable progress in understanding the cellular and connectional organization of the LFC, there are discrepancies in the reported maps. Some of the discrepancies stem from differences in the criteria employed to outline areas and/or the non-optimal sectioning of the gyrated primate cortex. Thus, differences between various maps (Vogt and Vogt, 1919; Walker, 1940; Barbas and Pandya, 1987; Barbas and Pandya, 1989; Petrides and Pandya, 1994; Matteli and Luppino, 2001; Petrides et al., 2005) give rise to controversies that need to be resolved. Such an endeavor is crucial since findings in the macaque LFC are indispensable because of the level of detail that they offer in generating hypotheses about the organization of the human LFC (e.g. Amiez and Petrides, 2009; Passingham and Wise, 2012; Margulies and Petrides, 2013). Specifically, with respect to the dorsal LFC, inconsistencies pertain to the presence of distinct cortical areas along the superior frontal region anterior to the end of the superior arcuate sulcus (Figure 1 A,C,E), the caudal premotor cortex (Figure 1 A,B,D,E), and the border of the dorsal and ventral motor/premotor areas (Figure 1 A,B,D). With respect to the ventral precentral area, namely the region that extends from the central sulcus to the region that surrounds the inferior ramus of the arcuate sulcus, several areas have been identified (e.g., Matelli et al., 1985; Barbas and Pandya, 1987; Petrides and Pandya, 1994; Petrides et al., 2005; Belmalih et al. 2009; Gerbella et al., 2007). There is, however, still debate concerning the extent and even the presence of certain cortical areas in this region in macaques. Such a debate and controversy obscures aspects concerning the evolution of areas and circuitry related to aspects of language. Specifically, the presence of a macaque homologue of part of the so-called Broca's region (area 44) in humans has been debated (Matelli et al., 2004). The presence of a cytoarchitectonic homologue of area 44 in the macaque inferior arcuate sulcus and its involvement with orofacial function has been established and clearly distinguished from ventral premotor areas (Petrides and Pandya, 1994; Petrides et al., 2005). This observation was recently confirmed in further cytoarchitectonic analysis of the ventrolateral frontal region in the macaque (Belmalih et al. 2009). Moreover, it has been shown that area 44 involved with orofacial/somatomotor functions lies at the fundus of the most ventral part of the inferior arcuate sulcus, while the cortex lying more dorsally is implicated in visuomotor attentional functions (area 8Av) (Petrides et al., 2005). It is desirable to obtain further evidence in order to corroborate and further elucidate the existence and borders of the distinct areas in the ventral precentral region.

**Figure 1.**
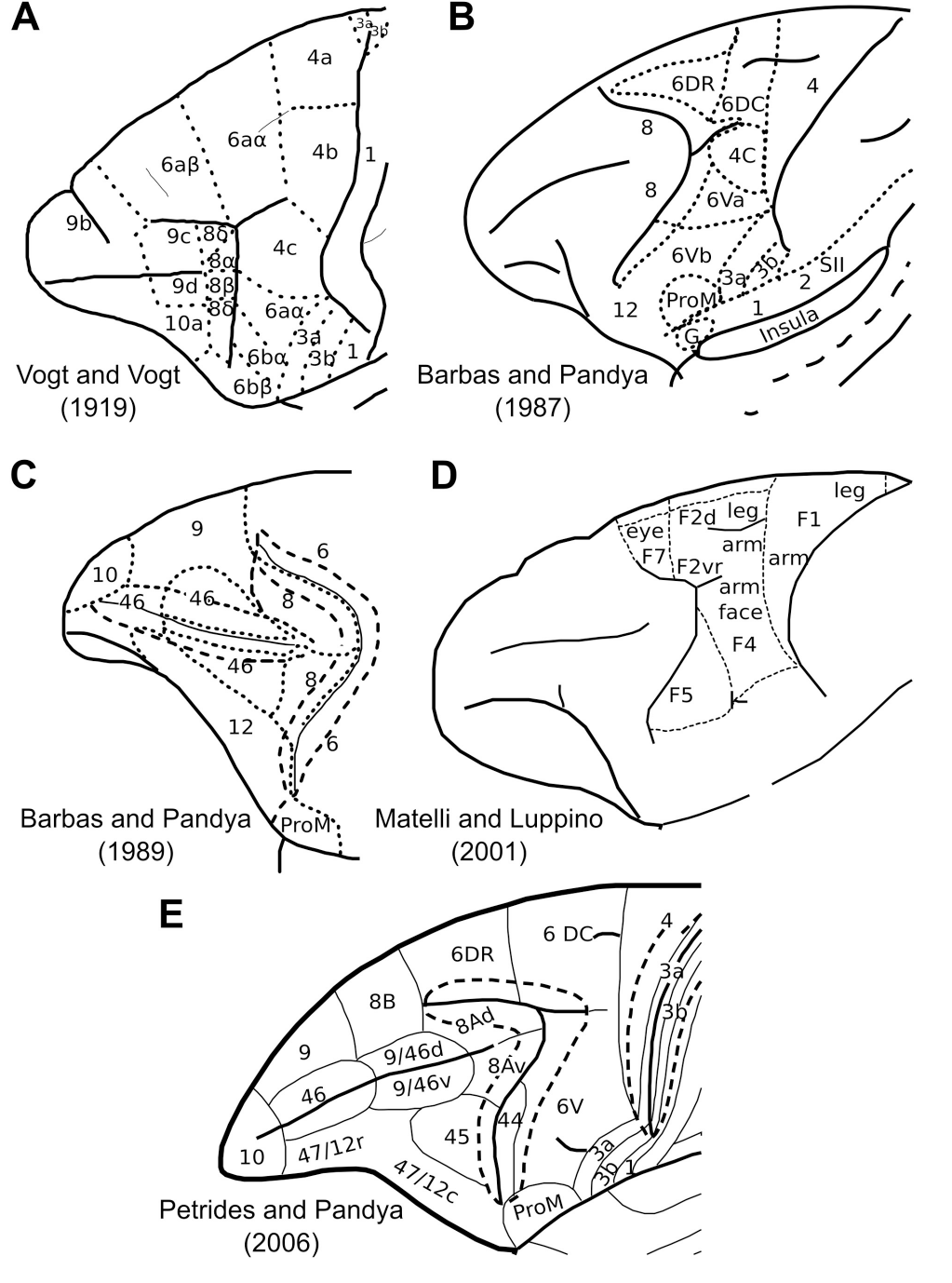
**A-E** Cytoarchitectonic maps of the lateral surface of the frontal cortex. Certain maps parcellate only parts of the lateral frontal surface.

In vivo neuroimaging techniques, such as resting-state fMRI (rsfMRI), can unveil distinct connectional division of the primate brain that are consistent, though not corresponding in a one-to-one fashion, with gold-standard tract-tracing findings (Margulies et al., 2009; Miranda-Dominguez et al., 2014). Because of their non-invasive nature such techniques are used for delineating areas in the human brain and demonstrate good co-localization with cytoarchitectonically defined areas (e.g. Kelly et al., 2010; Goulas et al., 2012; Margulies and Petrides, 2013).

In this study, we perform a data-driven connectivity-based parcellation of the LFC in the macaque monkey based on rsfMRI in order to inform controversies over existing organization schemes derived from invasive methods. Specifically, we aim to find evidence for connectional divisions and relate them to proposed parcellation schemes derived from histological analysis for which consensus is still lacking. RsfMRI, despite its disadvantage with respect to resolution and specificity, allows the connectivity-based parcellation of the whole extent of the LFC in a quantitative manner, whereas invasive tract-tracing techniques are restricted to a limited number of areas that can be injected. Clearly, rsfMRI is not a substitute of histological analysis but rather a complementary modality that can inform previous histologically-derived parcellation schemes. Lastly, a data-driven rsfMRI connectivity-based parcellation of the macaque LFC establishes the foundation for future macaque-human comparisons with the same modality (e.g, Margulies et al., 2009; Hutchison et al., 2012; Mantini et al., 2013; Salet et al., 2013) by overcoming the limitation of manual seed placement and adoption of specific a priori defined maps (Margulies et al., 2009; Salet et al., 2013).

## Materials and Methods

High-resolution rsfMRI data were acquired from 6 macaque monkeys at 7T. For each monkey, 10 runs of 150 EPI functional volumes (TR = 2000 ms; TE = 16 ms; flip angle=70°, matrix=96×96; FOV=96×96 mm; voxel size = 1 mm isotropic) were acquired, each run lasting 5 min. One T1-weighted anatomical image (TE = 2.5ms; TR = 2300ms; TI = 800ms; FOV = 96×96mm; 750p,m isotropic resolution) was also acquired (see Babapoor-Farrokhran et al., 2013 for details). Data were preprocessed with the REST toolbox (restfmri.net/forum/REST_V1.8) and SPM5 (Welcome Trust) and included realignment, slice-time correction, coregistration of functional and anatomical scans, regressing out white matter and cerebrospinal fluid signal, linear trends and six movement parameters. For the segmentation of the structural volumes, the macaque tissue priors provided in McLaren et al. (2009) were used. White matter and cerebrospinal fluid signal were extracted by using the corresponding probability tissue type images from each animal. A 0.8 threshold was applied to these images and subsequently the mean signal of the remaining voxels resulted in the white matter and cerebrospinal fluid regressors. In addition, band-pass filtering (0.01-0.1 Hz) and spatial smoothing (2mm FWHM) was applied. Such preprocessing steps are similar with those applied in previous rsfMRI macaque data (e.g. Hutchison et al., 2011; Sallet et al., 2013; Mantini et al., 2013).

The LFC was delineated on the F99 template (Van Essen, 2004) available in CARET (http://brainvis.wustl.edu/wiki/index.php/Caret:About) in order to create an LFC mask (Figure 2). We did not extend the posterior part of the mask until the fundus of the central sulcus, encompassing the presumed posterior limit of the primary motor cortex, in order to avoid examination of this region prone to partial volume effects and contamination of the fMRI signal between the posterior (somatosensory areas) and anterior (primary motor areas) banks of the central sulcus. The analysis was restricted to the left hemisphere for setting the foundation for subsequent comparative analysis with, presumed left lateralized, language-related areas/networks involving the human LFC. Moreover, we restricted the analysis to the left LFC for comparisons with histologically derived maps which mostly depict the left LFC. The LFC mask was transformed to the native space of each animal and the rsfMRI time courses of each grey matter voxel within the LFC patch were extracted. For each run, a within patch voxel-to-voxel correlation matrix was computed and these matrices were then averaged. The N×N, where N is the number of grey matter voxels within the LFC mask, average matrix from each animal was thresholded to result in a density of 0.01, thus creating a fully connected, undirected and weighted graph. Density is the ratio of connections/edges in the graph over the maximum possible edges in the graph given its number of nodes N (in our case number of voxels). A high sparsity, i.e. low density, for the graphs, which at the same time ensures full connectedness, was chosen in order to decrease computational time and detect modules/areas that otherwise might not be decipherable due to the “resolution limit” (Fortunato and Barthélemy, 2007) of the employed module detection algorithm. The Louvain module detection algorithm (Blondel et al., 2008) was applied as in Goulas et al. (2012) and incorporating the consensus strategy described in Lancichinetti and Fortunato (2012).

**Figure 2.**
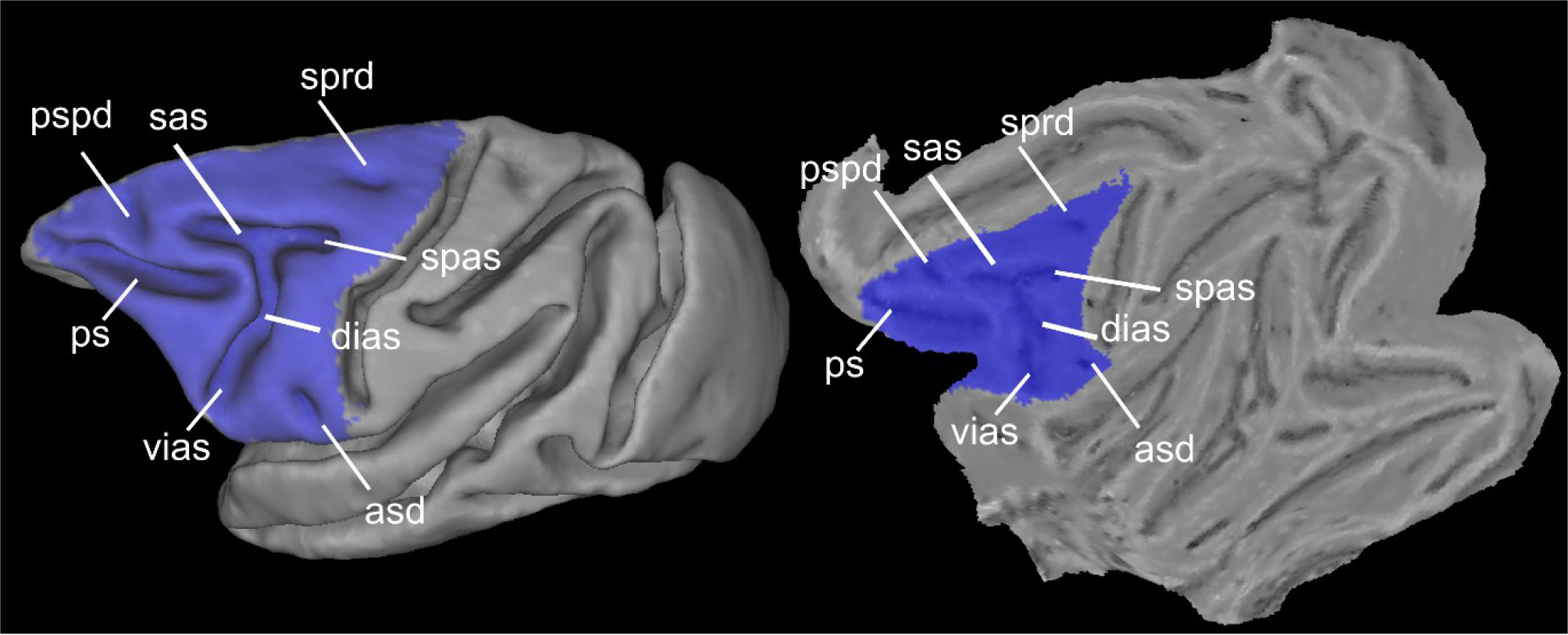
Spatial extend of the LFC mask used and major macroscopic landmarks within the LFC depicted on the F99 fiducial and flat surface. asd:anterior subcentral dimple; dias:dorsal compartment of the inferior branch of the arcuate sulcus; ps:principal sulcus; pspd:posterior supraprincipal dimple; sas:superior branch of the arcuate sulcus; spas:spur of the arcuate sulcus; sprd:superior precentral dimple; vias:ventral part of the inferior branch of the arcuate sulcus.

Briefly, the algorithm applies a greedy strategy for assigning each voxel to a module in order to maximize the modularity value Q (Blondel et al., 2008):

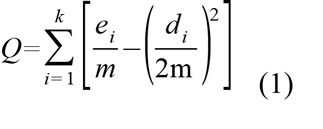

with *ei* representing the number of edges within module *i*, *di* representing total degree (i.e., number of functional connections/edges) of the nodes belonging to module *i*, and *m* representing the total number of edges in the graph. This value expresses how "surprising" the connectivity between voxels belonging to the same module is in relation to the connectivity expected by chance (see Blondel et al., 2008, for details). It should be noted that the number of modules are not determined a priori but derived from the algorithm and the dataset at hand. In other words, the number of modules is such so that the Q value is maximized. Moreover, the algorithm will always result in a solution and a corresponding Q value. Hence, the Q values obtained in the analysis are compared with what would be expected by chance by adopting two null models (see below). The aforementioned approach is stochastic, i.e. applying the algorithm many times does not guarantee the exact same solution/module decomposition. Moreover, these solutions might be substantially different and exhibit a high modularity value Q, a phenomenon termed "degeneracy of modularity" (Good et al., 2010). Because of the presence of equally good solutions, instead of picking up the solution with the highest value Q, the solutions can be combined with a consensus strategy (Lancichinetti and Fortunato, 2012). A N×N consensus matrix is formed from the solutions of the module detection algorithm, but now an entry i, j in the matrix does not denote the correlation of the rsfMRI time courses of voxel i and j but the frequency with which these voxels have been assigned to the same module. We adopted this consensus strategy because it is shown to lead to improved accuracy and stability of parcellation results (Lancichinetti and Fortunato, 2012). The above strategy has two free parameters, i.e. the number of solutions to form the consensus matrix and the threshold to be applied. We chose 100 as the number of solutions as input to the consensus clustering, since extensive previous analysis showed that above ~50 solutions there is a plateau in accuracies (see supplementary material in Lancichinetti and Fortunato, 2012). Moreover, the threshold parameter does not seem to influence the accuracy and consequently we chose a value of 0.5 to speed up the procedure (see supplementary material in Lancichinetti and Fortunato, 2012). The consensus matrix is then fed to the module detection algorithm after the application of the threshold (0.5), i.e. two voxels are assigned to the same module in half or more of the solutions, to produce 100 solutions. Subsequently, the consensus matrix is formed anew and the procedure is iteratively applied until the consensus matrix becomes a block diagonal matrix with ones (zeros) denoting voxels always assigned to the same (different) module.

The "resolution limit" of the module detection algorithm (Fortunato and Barthélemy, 2007) can lead to the merging of distinct modules (in our case distinct LFC areas). Thus, informed by previous cytoarchitectonic parcellation schemes, potentially merged modules will be taken into account separately and fed into a second parcellation. This approach is suggested for investigating further subdivisions that may be concealed in the results of the first parcellation (Fortunato and Barthélemy, 2007; Ruan and Zhang, 2008) and has been previously applied in neuroimaging analysis (Nelson et al., 2010).

In order to assess the statistical significance of the parcellation resulting from the module decomposition, two null models were adopted, i.e. the degree-preserving rewiring null model (Rao and Bandyopadhyay, 1996; Maslov and Sneppen, 2002), with degree in our case denoting the number of functional connections of a voxel, and the null correlation matrix model (Zalesky et al.,2012). While the former model preserves certain topological properties of the original network, i.e. the degree distribution, the latter aims at creating null correlation matrices that preserve the distribution of the correlation values of the original network and the increased clustering of the network introduced by the correlation itself (Zalesky et al., 2012). Briefly, the degree-preserving rewiring null model is derived as follows:Two pairs of interconnected nodes (a-b, c-d) are randomly selected and rewired be swapping partners, i.e. a-d, b-c. The process is repeated many times, here 100, so that any topological pattern of the original network, apart from degree-distribution, number of nodes and edges, is destroyed. The null correlation matrices were generated with the Hirschberger-Qi-Steuer algorithm that creates correlation matrices with matched mean and variance to the original matrices (see for details the Appendix in Zalesky et al., 2012).

For assessing the stability of the parcellation results, the aforementioned analysis was conducted in the odd and even runs separately. Hence, for each animal two partitions derived from the odd and even runs were obtained. The more similar these partitions are with the partition obtained with all the runs and in between them, the more stable the solutions can be considered. The similarity of the partitions was quantified with the normalized variation of information (Meila, 2007). This metric has theoretical values in the range [0,1], with 0 indicating identical partitions and 1 completely different.

The above approach resulted in a module map for each animal that can be considered to correspond to distinct areas. The module maps from each animal were grouped together in a data-driven manner after normalization to F99 space by using the center of mass as a similarity criterion (Goulas et al., 2012). This resulted in a probability map for each module denoting in each voxel the frequency of colocalization of each module across the animals. For estimating the functional connectivity (FC) map of each module at the group level, a spherical seed (1.5 mm radius) was placed at the weighted center of mass of each probability map. To ensure the placement of the seed in the most “representative” coordinate, before the calculation of the weighted center of mass, the probability maps were thresholded in order to contain voxels denoting colocalization in at least two animals. Time series from the seeds were extracted and entered as regressors in a multiple regression model combining all runs from all animals in a fixed effects analysis. This procedure resulted in FC maps for each module at the group level. The maps were thresholded at a cluster-level q<0.05 (FWE) (cluster defining threshold:p<0.001 uncorrected). In order to quantify the similarity of the FC maps, all pairwise spatial similarities were computed by calculating the 1 − r between the coefficients ('con*.nii" maps in SPM) for each module derived from the aforementioned fixed effects analysis. For computing pairwise similarities we restricted the analysis in grey matter voxels after segmenting the F99 template. The pairwise similarities were used to construct a dendrogram with the average linkage method. The faithfulness of preservance of the original distances in the dendrogram was assessed with the cophenetic coefficient.

The analysis was performed with custom Matlab (The Mathworks) functions and functions from the Brain Connectivity Toolbox (Rubinov and Sporns, 2010).

## Results

The module detection algorithm resulted in high modularity values (Q mean:0.83, std:0.01, p<0.01) compared with those obtained from both null models (Q_rewned_ mean:0.26, std:0.04, Q_null correlation_ mean:0.27, std:0.04). Good correspondence between the partitions estimated separately from the odd and even runs was observed resulting in very low variation of information, and thus highly similar partitions (mean:0.13, std:0.02). The observed similarity of the partitions from the odd and even runs was much higher when compared to the randomized affiliation vectors derived from the module detection algorithm (mean:0.65 std:0.03). In addition, the variation of information in the partitions obtained from the odd and even runs were very similar with the one obtained from all runs (variation of information mean:0.09, std:0.02). These results highlight that our parcellations are characterized by stability, in the sense that the similarity of partitions obtained from odd and even runs give rise to values very near the theoretical maximum similarity and substantially differ from values expected by chance, and statistical significance, since the Q values differed from the ones obtained from the two null models.

The modules obtained in each animal resulted in 14 clusters of modules, hereafter clusters, at the group level (Figure 3 A). All these clusters were formed from modules that were present in at least 5/6 animals. The spatial layout of the clusters indicates a neuroanatomically realistic parcellation (see below and Figure 1). Certain clusters seem to encompass more than two distinct cortical areas (Figure 3 A with outlined borders). More specifically, the green outlined cluster (Figure 3 A) seems to encompass areas 6aα and 4b of Vogt and Vogt (1919) (Figure 1 A). The brown outlined cluster (Figure 3 A) seems to encompass areas ProM, 3a, 3b and 1 (Figure 1 E). The blue outlined cluster (Figure 3 A) seems to encompass areas 45 and 44 (Figure 1 E). The purple outlined cluster (Figure 3 A) seems to encompass areas 47 and parts of 9/46v (Figure 1 E). Lastly, the light purple outlined cluster (Figure 3 A) seems to encompass areas 46, 9 and 10 (Figure 1 E). To find out if these clusters could be further subdivided, they were submitted to a separate second pass parcellation. This resulted in parcellations with higher than chance Q values (mean:0.80, std:0.01, p<0.01, Q_rewired_ mean:0.29, std:0.05, Q_null correlation_ mean:0.48, std:0.08, p<0.01). Again, the modules were grouped across the animals resulting in 10 clusters. These clusters, along with the ones from the first pass analysis resulted in 19 clusters (C1-C19) (Figure 3 B). This parcellation is the focus of our subsequent analysis. Specifically only the clusters located at the dorsal LFC, i.e. C1-C8, and ventral LFC, i.e. C14-C19 (Figure 3 B), will be the focus of the present report. The results from the clustering in the principal sulcus were not interpretable in terms of prior parcellation schemes and therefore not satisfactory. This is presumably due to the partial volume effects and signal contamination of the fundus, dorsal and ventral banks of the principal sulcus, rendering the uncovering of the connectional heterogeneity of this LFC region problematic. Higher resolution data seem necessary for examining this region with rsfMRI (see also Limitations and future directions).

**Figure 3.**
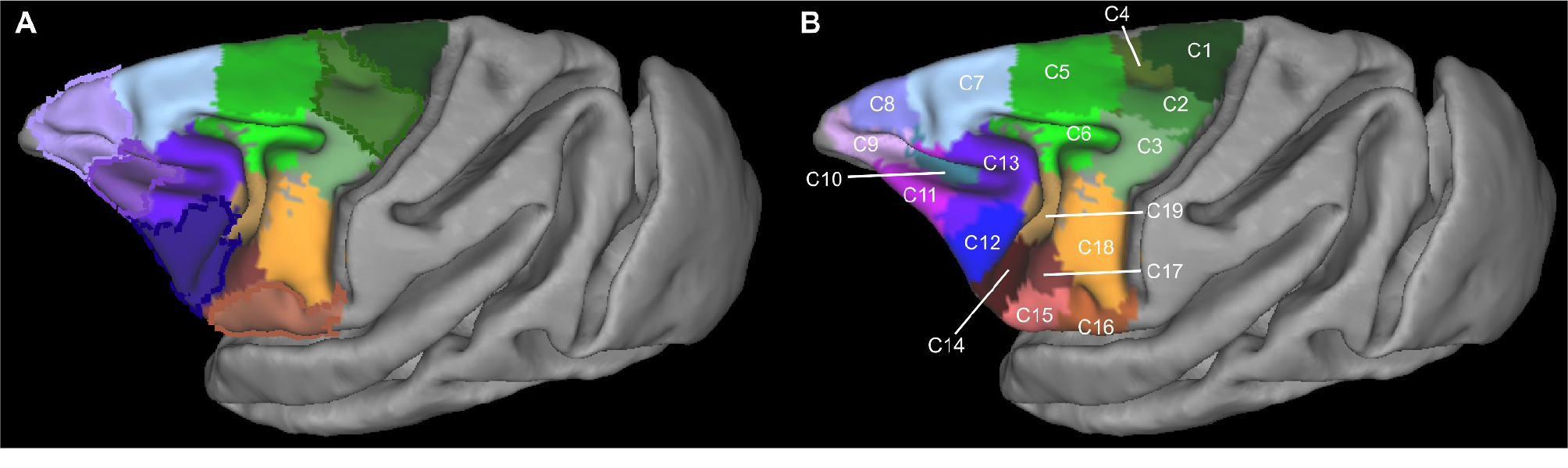
Summary of the parcellation results of the first (**A**)and second (**B**) “pass” of the algorithm. The depicted results constitute winner-takes-all maps. The modules constituting each cluster are forming a probabilistic map (see Figure 4). These probabilistic maps are combined to produce the depicted winner-takes-all maps by assigning each voxel a unique integer corresponding to the cluster exhibiting the highest probability in this voxel. Subsequently, each cluster is coded with a unique color and named arbitrarily as C1, C2,… C19. This cluster-wise colour scheme is also followed in Figures 4 and 5. Spatial location of the clusters dictates their colour ‘family':dorsal motor/premotor clusters are colour coded with shades of green, ventral premotor clusters with brown/orange, and the prearcuate ones with blue/violet. Borders around clusters in **A** indicate the ones that were further subdivided on the basis of the second pass results depicted in **B** (see Results).

The FC maps of each cluster that were fed into a hierarchical clustering resulted in the grouping of the clusters into three broad connectivity families, a dorsal motor/premotor, a ventral premotor and a prearcuate/prefrontal (Figure 4). The dendrogram was constructed with the average linkage method since this method resulted in the most faithful representation of the original distances, as assessed with the cophenetic coefficient (0.74), when compared to single (0.51), complete (0.67), and weighted (0.72) linkage methods. A notable exception in the connectional segregation, which largely coincides with the spatial segregation of the clusters, was postarcuate C6 that was not grouped with the dorsal motor/premotor clusters but with the prearcuate ones (Figure 3 B, Figure 4).

**Figure 4.**
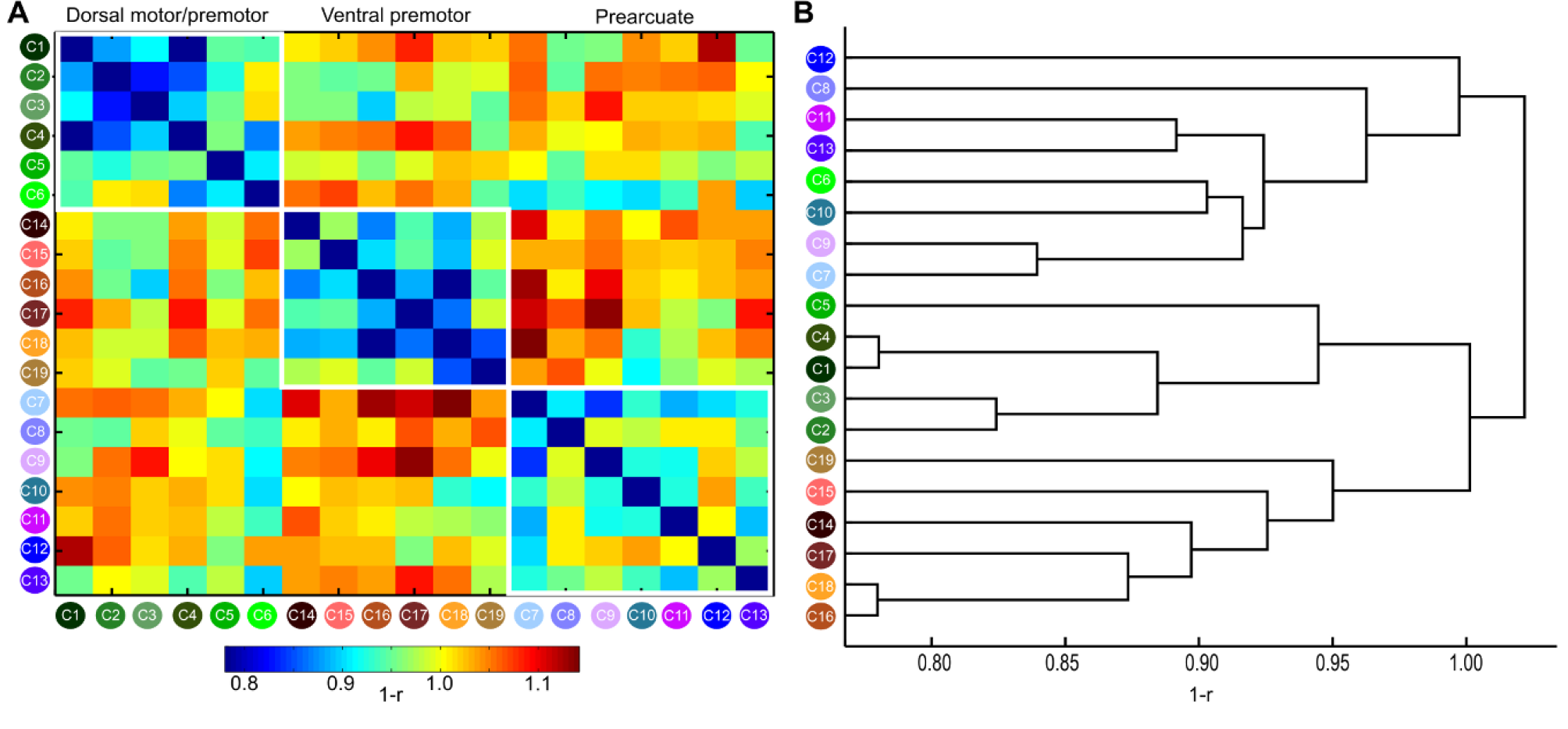
**A** Connectivity similarity matrix for all the clusters. Note that the clusters are arranged based on their spatial location to three broad groups:dorsal and ventral motor/premotor and prearcuate indicated by the white outlines. Connectivity similarity was measured as 1-r, where r Pearson's correlation coefficient hence lower (higher) values indicate higher (lower) connectivity similarity (see Materials and Methods). **B**, Dendrogram constructed based on the connectivity similarity of the clusters (see Material and Methods and Results). Three broad groups are discernible,broadly coinciding with the groups defined by spatial location (Figure 3B). Thus, groups discernible based on spatial or connectional information largely coincide (compare the groupings of the clusters in **A** and **B**). A notable exception is C6 which seems more affiliated, on a connectional basis, with the prearcuate group, despite its postarcuate location (see Results and Discussion).

Below we document the results and interpret the dorsal and ventral LFC clusters on the basis of histologically defined cortical areas based on topographic and FC information. A qualitative comparison is necessitated by the lack of quantitative probabilistic maps in a stereotaxic space for the macaque LFC. We first describe the dorsal LFC results proceeding along the dorsal part of the frontal lobe following a caudal-rostral direction from the central sulcus to the frontal pole, followed by the results on the ventral LFC.

### Dorsal LFC

#### Cluster C1

In the most dorso-caudal part of the frontal lobe, immediately in front of the central sulcus, there is cluster C1 which most probably corresponds, from a topographic perspective, to a subdivision of the primary motor cortex, defined as area 4a by Vogt and Vogt (1919) (Figure 1 A). Its weighted center of mass (WCOM) in F99 space is (x=−7.4 y=−10.0 z=24.3) (All subsequent coordinates are in F99 space) (Figure 5). On the basis of electrical stimulation data, this region of the motor cortex corresponds to the trunk and lower limbs of the body (Vogt and Vogt 1919; Woolsey, 1952). C1 is characterized by connectivity with the medial wall of the primary motor cortex, the adjacent supplementary motor cortex, and the caudal cingulate motor areas (Picard and Strick, 2001). There was also strong connectivity with the superior parietal lobule (areas PE and PEc) and anterior part of the intraparietal sulcus (Petrides and Pandya, 1984; case 1 in Bakola et al., 2013). This region has connectivity with the adjacent part of the primary motor cortex (area 4b) and the rostrally adjacent dorsal premotor cortex (F2/6DC). Connectivity was restricted to the dorsal motor and premotor areas (Figure 6), consistent with the presumed evolutionary origins of these areas from the archicortical trend (Barbas and Pandya, 1987). This affiliation with the dorsal constellation of LFC areas was also evident when quantifying the similarity of the whole brain connectivity with the rest of the clusters, since C1 belongs to the dorsal motor/premotor group (Figure 4).

**Figure 5.**
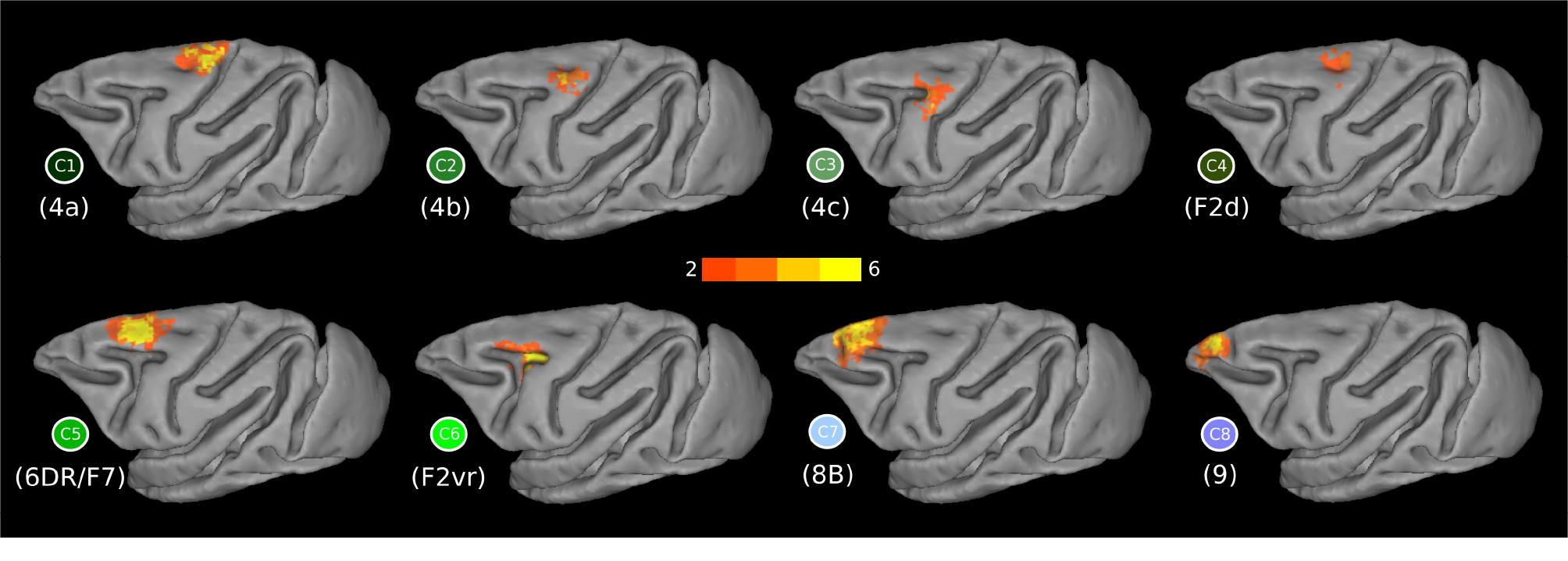
**Figure 5.** Probabilistic maps of the dorsal LFC clusters and their respective functional connectivity maps. Orange to yellow colours in the probabilistic maps denote low to high overlap across animals. The faded coloured borders correspond to the dorsal clusters as depicted in the winner-takes-all map in Figure 3B.

**Figure 6.**
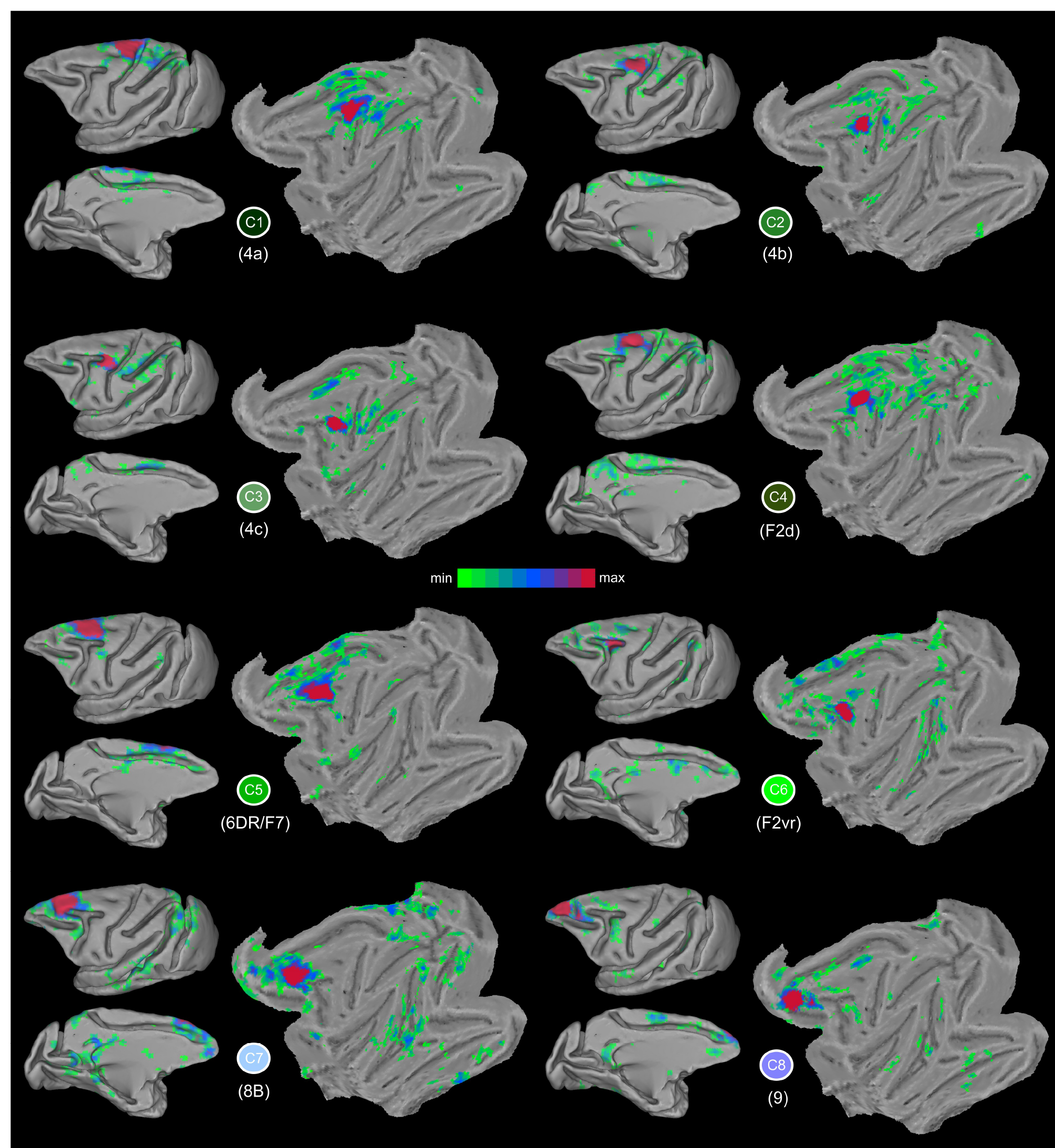
Functional connectivity maps of the dorsal clusters in the F99 template. Higher t values are denoted with red/purple colours. Each cluster is denoted by its unique colour (see Figure 3B).

#### Cluster C2

C2 is located dorso-caudaly to the most posterior part of the spur of the arcuate sulcus and ventral to C1 (WCOM x=−13.4 y=−6.1 z=21.7) (Figure 5). Its location corresponds well with the 4b subdivision of primary motor area 4 by Vogt and Vogt (1919) (Figure 1 A) corresponding to the forearm, finger and shoulder representations. Its connectivity pattern in the medial wall involves SMA (cases 3 and 7 in Morecraft and Van Hoesen, 1993) and area PGm (case 9 in Petrides and Pandya, 1984). On the lateral surface, it involves areas PE (case 1 in Bakola et al., 2013) and its extension to the intraparietal sulcus, i.e. area PEa (case 1 in Petrides and Pandya, 1984) (Figure 6).

#### Cluster C3

C3 is located posterior to the spur of the arcuate sulcus, with a focus on the posterior-most part of the spur (WCOM x=−16.6 y=−1.9 z=18.6) (Figure 5), possibly involving the part of the primary motor cortex that Vogt and Vogt (1919) refered to as 4c (Figure 1 A) eliciting facial and neck responses. This region is the focus of connectivity from the anterior part of the superior parietal lobule and the anterior part of the adjacent medial bank of the intraparietal cortex (case 1 in Petrides and Pandya, 1984). The connectivity of C3 is prominent with the pre-SMA on the medial wall, as well as parietal area PGm. Extensive connectivity was also observed with the rostral part of the intraparietal sulcus and the rostral superior parietal lobule (area PE) (case 2 in Bakola et al., 2013) (Figure 6). The connectivity of C3 is clearly affiliated with the dorsal motor/premotor group (Figure 4).

#### Cluster C4

C4 (WCOM x=−8.2 y=−4.9 z=23.4), which lies anteroventral to C1, is focused around the superior precentral dimple (Figure 5), where area 6DC (also known as F2) is located. More specifically, C4 is colocalizing with dorsal subdivision F2 (Luppino et al, 2003) (Figure 1 D). The connectivity of C4, consistent with tract-tracing studies, is with supplementary motor cortex and to a lesser extent with the cingulate motor areas (Luppino et al., 2003). In addition, it exhibits strong connectivity with the superior parietal lobule (areas PE and PEc), the adjacent intraparietal sulcus (Petrides and Pandya, 1984; Marconi et al., 2001), the inferior parietal lobule (case 1 in Petrides and Pandya, 1999) and the medial parietal region, especially area 31, and the more dorsal anterior area PEci (case 13 in Morecraft et al., 2012) (Figure 6). C4 connectivity assigns this cluster to the dorsal motor/premotor group (Figure 4).

#### Cluster C5

Another distinct cluster, C5 (WCOM x=−8.4 y=4.1 z=21.8), is located above the superior branch of the arcuate sulcus, which corresponds to the location of area 6DR (also known as F7) (Figure 1 D, E and Figure 3). The peak of the probabilistic map lies in the anterior part of this cluster (Figure 5). Similar to C4, C5 shows strong connectivity with the adjacent dorsal premotor cortex and the adjacent medial wall of the frontal lobe where the pre-SMA region lies. The connectivity also extends into the cingulate sulcus involving the cingulate motor areas. This pattern is consistent with tract-tracing results (case 13 FB in Luppino et al. 2003). C5 is distinguishable from C4 in its more anterior connectivity along the medial wall to the pre-SMA, as opposed to C4 connectivity with the more posteriorly located SMA (Luppino et al, 2003) (Figure 6). C5 connectivity assigns this cluster to the dorsal motor/premotor group (Figure 4).

#### Cluster C6

C6 (WCOM x=−12.6 y=2.8 z=16.4) is focused around the spur of the arcuate sulcus with the peak of the probability map in the posterior part of the spur (Figure 5). The topography resembles the subdivision of F2 described as F2vr by Luppino et al. (2003) (Figure 1 D) (see also Discussion). Consistent with tract-tracing studies involving area F2vr, the C6 connectivity pattern involves the cingulate motor areas, i.e. CMAd, CMAv, CMAr, and parts of dorsal prefrontal cortex (Luppino et al., 2003). In addition, there was connectivity with the dorsal prelunate region, the parieto-occipital sulcus (case 2 in Yeterian and Pandya, 2010; Stepniewska et al., 2005), the vicinity around the accessory parieto-occipital sulcus possibly hosting areas V6/V6A (Luppino et al., 2005), the medial intraparietal area (Marconi et al., 2001), and the caudal superior parietal lobule (case 6 in Petrides and Pandya, 1984) (Figure 6). Interestingly, despite the fact that C6 lies partly within the caudal bank of the arcuate sulcus (Figure 5), its connectivity pattern is clearly more similar to prearcuate clusters when compared to the dorsal motor/premotor ones (Figure 4).

#### Clusters C7 and C8

Along the superior frontal region, anterior to C5, two distinct clusters, C7 (WCOM x=−7.8 y=14.3 z=18.7) and C8 (WCOM x=−6.5 y=22.4 z=14.8), were uncovered (Figure 5). In the past, this region had been treated as either two distinct areas, namely areas 8B and 9 by Walker (1940) and Petrides and Pandya (1994), or as one, i.e. area 9 (Barbas and Pandya, 1989) (Figure 1 C, E). Our results demonstrate that, on a connectional basis, two distinct clusters could be distinguished.

C7 extends from the anterior end of the superior branch of the arcuate sulcus and continues as far as the posterior supraprincipal dimple, which is the region were area 8B lies (Figure 1 E). This area marks the transition from premotor areas to the prefrontal region, as evident in the shift of the connectivity profile from C5 to C7 (Figures 4 and 6). Anterior to the posterior supraprincipal dimple lies C8, which corresponds to the location of area 9 as defined by Walker (1940) and Petrides and Pandya (1994) (Figure 1 E). Both clusters are strongly connected with the retrosplenial cortex, a characteristic of the dorsolateral prefrontal cortex (Morris et al., 1999). The subtle difference in retrosplenial connectivity is noted in C7, putative area 8B, as being slightly more anteriorly focused, whereas C8, putative area 9, is more ventral in retrosplenial cortex (Figure 6) (see figures 7 and 10 in Morris et al., 1999; case 5 in Petrides and Pandya, 1999; cases 3 and 4 in Pandya and Yeterian, 1996). In addition, C7 demonstrates connectivity to area Opt (case 2 in Petrides and Pandya, 1999) a pattern that is not pronounced for the more anterior C8. The connectivity of C8 with the ventral premotor cortex does not seem at odds with results from tract-tracing studies and most likely arises due to polysynaptic/network effects (Figure 6) (see Discussion).

### Ventral LFC

All of the clusters documented below were assigned to the ventral premotor group (Figure 4).

#### Cluster C14

This cluster was located at the fundus of the ventral part of the inferior arcuate sulcus (vias) (Figures 1 E, 2, 7) (WCOM x=−20.9 y=9.0 z=5.3) where area 44 has been identified (Petrides and Pandya, 1994; Petrides et al., 2005), an area distinct from the posteriorly adjacent ventral premotor clusters C15 and C17 and the anteriorly adjacent prearcuate cluster C12. More recent histological analysis has confirmed the presence of a distinct area 44 in the fundus of the inferior arcuate sulcus that differs from posterior premotor area F5 (Belmalih et al., 2009). We conclude that C14 co-localizes very well with what has been identified as area 44 in independent histological analyses of the ventral part of the inferior arcuate sulcus.

The FC of this region was characterized by strong links to area PFG in the anterior part of the inferior parietal lobule and the adjacent intraparietal cortex, often referred to as area AIP, which is consistent with prior findings from gold-standard tract-tracing studies (case 2 in Petrides and Pandya, 2009; cases 1 and 3 in Frey et al., 2014). On the medial part of the hemisphere, FC was observed with the cingulate motor areas and more anterior portions of the cingulate cortex extending around the genu of the corpus callosum. In addition, FC was observed with the insular cortex and the secondary somatosensory region (cases 1 and 2 in Frey et al., 2014) (Figure 8). A noteworthy discrepancy with knowledge from invasive tract-tracing studies is the lack of FC with the ventral lip of the principal sulcus hosting area 9/46v.

#### Cluster C15

Posterior to C14 and immediately anterior to the anterior subcentral dimple (asd), a distinct cluster was uncovered, i.e. C15 (Figure 7) (WCOM x=−25.2 y=6.4 z=3.9). The extent of the cluster was bounded by the imaginary line at the dorsal part of the asd and largely avoided the posterior bank of the vias (Figure 7). From a topographic point of view, the cluster appears to correspond to area ProM, namely the proisocortical motor cortex (Sanides, 1968; Barbas and Pandya, 1987). This area is considered as the proisocortical architectonic step from which subsequent differentiation led to the ventral premotor areas (Barbas and Pandya, 1987).

**Figure 7.**
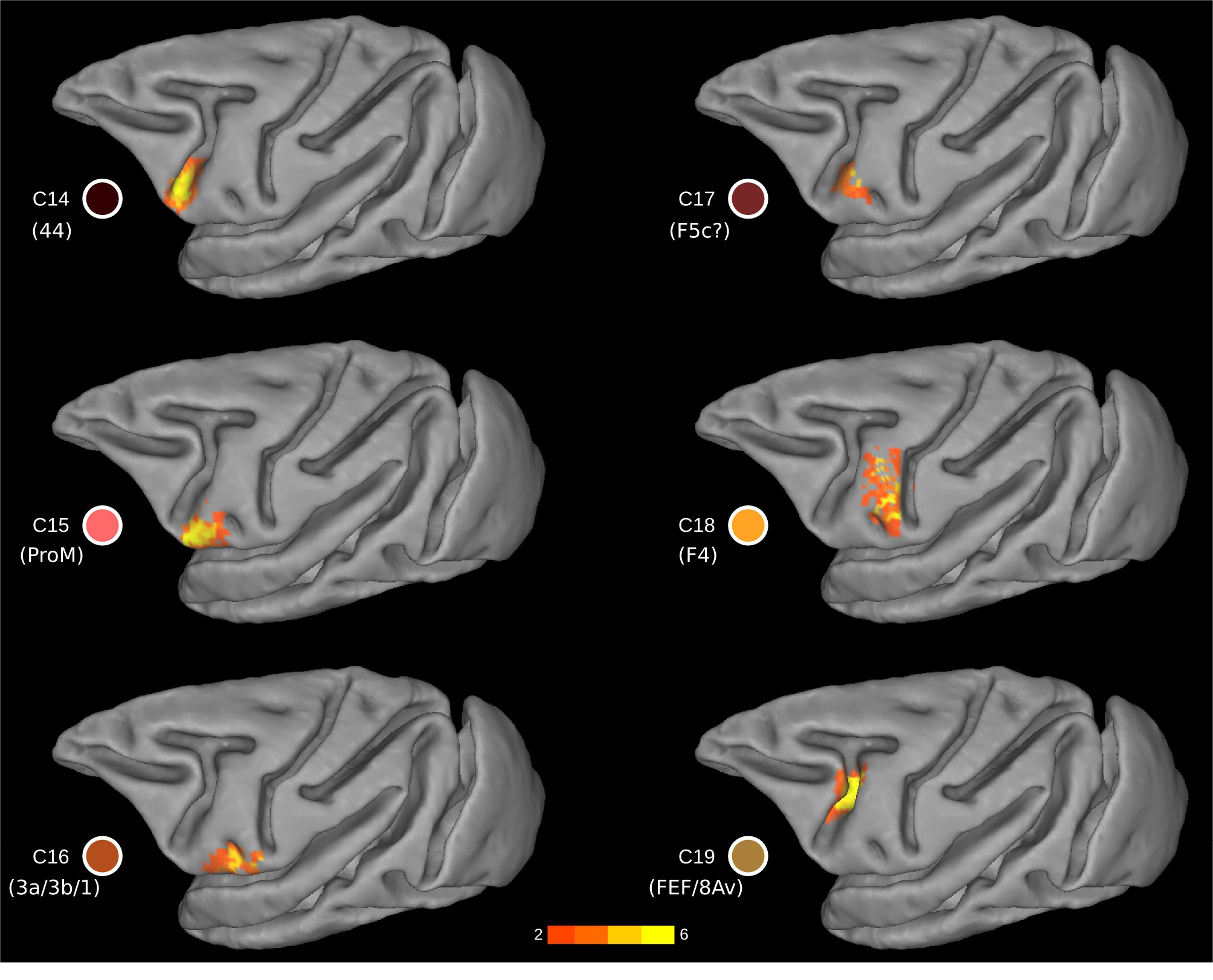
Probability maps for each ventral LFC cluster. The probability maps are thresholded in order to include part of the cluster present in at least two animals. The colour of the clusters corresponds to the colour-coding scheme of Figure 3B. Yellow (orange) colours denote high (low) probability, i.e. presence across animals.

The FC of C15 was mostly local (Figure 8). There was FC with 6VR, 6VC, and the nearby opercular zone including the most anterior part of the insula and also the secondary somatosensory region. The FC seemed to extend into the most ventral part of the central sulcus, possibly involving the orofacial part of the somatosensory region (ARG case 1 in Cipolloni and Pandya, 1999). On the medial wall, FC possibly corresponding with area SMA was observed (FRT case 1 in Cipolloni and Pandya, 1999) (Figure 8).

**Figure 8.**
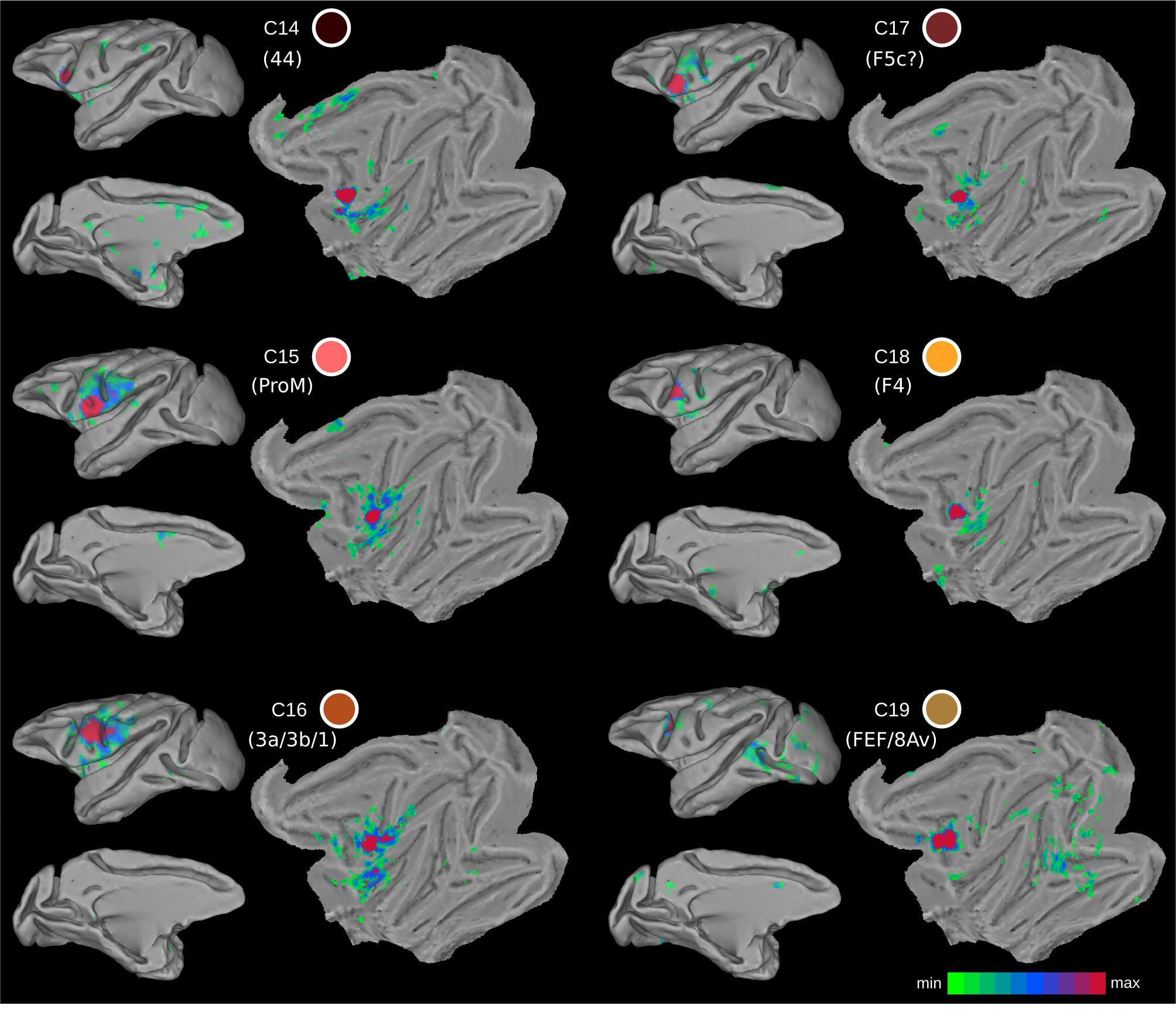
Functional connectivity maps of the ventral LFC clusters. The colour of the clusters corresponds to the colour-coding scheme of Figure 3B. Shades of red in the functional connectivity maps denote higher t values.

#### Cluster C16

In the most ventral part of the precentral region, in front of the ventral tip of the central sulcus and posterior to the asd (WCOM x=−26.4 y=1.2 z=4.6), a cluster is located that appears to correspond with the precentral extension of the primary somatosensory region (areas 3 a, 3b, 1), as first described by Vogt and Vogt (1919) (Figures 1 and 7). Somatosensory cortical areas 3, 1, 2 are primarily found on the postcentral gyrus of the macaque monkey brain, but continue around the most ventral part of the central sulcus and occupy a part of the precentral gyrus as far as the asd. Cluster C16 is consistent with available architectonic maps of the macaque monkey frontal cortex that place this somatosensory region posterior to the asd, while proM extends anterior to this sulcus (Figure 1).

The local FC of C16 was with nearby precentral ventral somatomotor areas, including the anterior insula, the ventral part of the postcentral gyrus involving areas 3, 1 and 2 (cases 1, 2, 3 in Cipolloni and Pandya, 1999). Moreover, the FC pattern included the orofacial part of area 4, possibly 6VC, and anterior insula and nearby opercular areas and possibly including the ventral portion of 6VR. Overall, the FC pattern is restricted to ventral precentral and postcentral regions (Figure 8).

#### Cluster C17

This cluster was located on the postarcuate convexity and in the posterior bank of the ventral ramus of the inferior arcuate sulcus (Figures 1 B and 7) (WCOM x=−23.1 y=5.8 z=8.4). From a topological point of view, it is reminiscent of area F5 (Matelli et al., 1985) and colocalizes with F5c (Belmalih et al., 2009). The FC pattern of the cluster was predominantly local, including the anterior insular cortex and the ventral orofacial parts of the primary and secondary somatosensory cortex. These connections are consistent with those reported for the larynx area of the ventral premotor region (Simonyan and Jürgens, 2005) (Figure 8). In addition, sparse FC was observed with putative SMA in the medial wall and putative area PF in the parietal cortex, in line with invasive tract-tracing data (see case 36l and 42l in Gerbella et al., 2011). There are however certain noteworthy discrepancies. There was a lack of FC with areas of the granular frontal cortex, contrary to evidence from invasive studies (see case 36l and 42l in Gerbella et al., 2011).

#### Cluster 18

A separate cluster, C18, was observed dorsal to the ads, occupying the ventral part of the precentral region (Figures 1 B and 7) (WCOM x=−24.1 y=0.7 z=11.5). Its topography matches well with the ventral premotor area F4 (Matteli et al., 1985) with the exception that it does not extend dorsally until the spur of the arcuate sulcus and may also include a small part of the dorsal part of premotor area F5. The region around the spur of the arcuate sulcus appears as a distinct cluster (C3), corresponding to the region identified as area 4C by Vogt and Vogt (1919) and Barbas and Pandya (1987). The dorsal border of C18 appears to be the imaginary posterior extension of the principal sulcus (see Discussion).

The FC pattern of cluster C18 involves the rostral inferior parietal lobule, possibly area PF, and the rostral intraparietal sulcus, possibly area VIP, consistent with tract-tracing results (Luppino et al., 1999; Rozzi et al., 2006) (Figure 8). Area PF is mostly somatosensory related, whereas area VIP seems to include visual and tactile neurons (Geyer et al., 2012). The parieto-frontal circuitry formed by F4/F5 and VIP has been suggested to be functionally involved in the execution of movements for reaching and grasping objects in the environment (Geyer et al., 2012).

#### Cluster C19

Dorsal to C14 and still within the fundus of the arcuate sulcus, there was a distinct cluster that occupied the dorsal compartment of the inferior arcuate sulcus (dias) (WCOM x=−17.4 y=4.8 z= 11.1). At this level of the inferior arcuate (i.e. posterior to the end of the principal sulcus) lies cortex implicated in oculomotor control (the frontal eye field (FEF) region) (Huerta et al., 1987; Koyama et al., 2004; Bruce and Goldberg, 1984; Petrides et al., 2005) (Figures 1 B and 7). However, C19 might also encompass parts of the ventral premotor areas since it also encompasses parts of the posterior bank of the arcuate sulcus.

The FC of this cluster is consistent with reports from invasive methods. Strong FC was observed with the intraparietal sulcus and the nearby dorsal prelunate gyrus (area V4) (Ungerleider et al., 2008; Huerta et al., 1987; case 3 from Petrides and Pandya, 1999). In addition, the occipitotemporal transition zone close to the superior temporal sulcus, where area MT lies, was also part of the FC signature of this cluster (Huerta et al., 1987). Moreover, weak connectivity with area V1 is demonstrated in an invasive tract-tracing study (Markov et al., 2014), which potentially accounts for the observed V1 FC in our results (Figure 8). It should also be noted that similar functional properties with the FEF seem to characterize the cortex near the genu of the arcuate sulcus (C13 in Figure 3 B), possibly hosting head movement and large amplitude saccade related neurons (Zinke et al., 2015).

## Discussion

We have parcellated the LFC of the macaque based on rsfMRI. The resulting clusters, both in terms of their topography and connectivity, are consistent with several aspects of previous parcellation schemes based on cytoarchitectonic analysis (Figures 1, 3, 4 and 5). The current organization scheme based on the intrinsic functional architecture of the macaque LFC provides information relevant to certain debates on LFC organization. We elaborate on these aspects in detail below.

### Dorso-caudal premotor cortex (area 6DC/F2):The cortex in the superior precentral dimple and the cortex within the spur of the arcuate sulcus constitute distinct connectional divisions

The caudal part of the dorsal premotor cortex is an agranular cytoarchitectonic region that has been referred to as area 6aα by Vogt and Vogt (1919), as area 6DC by Barbas and Pandya (1987), and as area F2 by Matelli et al. (1985). Connectional and functional data suggest an orderly arrangement of somatomotor inputs related to the leg and arm more dorsally near the superior precentral dimple. The ventral limit of this region, however, has been problematic. Findings suggest distinct connectional and functional features of the cortical region within and near the spur of the arcuate sulcus. For instance, the cortex within the spur of the arcuate sulcus is strongly connected with area 45 (case 2 in Petrides and Pandya, 2002) and area 8Ad (case 5 in Petrides and Pandya, 1999), but neither area 45 nor area 8Ad connects with any part of the cortex dorsal to the spur in area 6DC. In other words, the multisensory prefrontal area 45 and the visuo-auditory prefrontal area 8Ad connect with the cortex of the spur but not with the cortex dorsal to the spur which receives massive input from somatomotor areas, such as PE (see Petrides and Pandya, 1984). Some of these connectional features are reflected in the FC of C4 and C6 (Figure 6). Furthermore, the cortex within the spur of the arcuate sulcus participates in oculomotor (Koyama et al., 2004) and visuomotor functions (Marconi et al., 2001; Fogassi et al., 1999) suggesting that this part of the cortex is a distinct area. Luppino et al. 2003 have referred to an area just above the spur as F2vr and this may partly overlap with cluster C6, although C6 lies primarily within the spur and extends slightly above and below it (Figure 3 B). There is also some immunohistochemical evidence consistent with the aforementioned subdivisions (Geyer et al., 2000).

The present resting-state functional connectivity analysis provides clear evidence that the cortical region within the spur of the arcuate sulcus is a distinct area, likely corresponding to F2vr (C6 in Figure 3 B). This area is clearly differentiated from other dorsal premotor areas, i.e. cluster C5, corresponding to area 6DR and cluster C4, corresponding to the dorsal part of area 6DC/F2 (Figure 3 B). These three dorsal premotor divisions exhibit very distinct connectivity profiles (Figures 6). Notably, C6/F2vr, despite the fact that, from a topographic point of view it is a postarcuate cluster, on a connectional basis, it resembles more clusters of the prearcuate group (Figures 4 and 6). In conclusion, there are at least two distinct areas discernible on a connectional basis within what has been previously defined as F2/6DC.

### The intrinsic resting state connectivity distinguishes two areas in the superior prefrontal region,consistent with areas 9 and 8B and places their border along the posterior supraprincipal dimple

The superior frontal region of the monkey, along the midline and anterior to the superior arcuate sulcus, has been considered as a single cytoarchitectonic area, labeled area 9 in some architectonic maps (Brodmann, 1905; Vogt and Vogt, 1919; Barbas and Pandya, 1989), while other maps have considered the caudal part of this region to be a separate area, labeled as area 8B (Walker, 1940; Petrides and Pandya, 1994) (Figure 1 A,C,E). Thus, ambiguity still characterizes the organization of the superior frontal region. The present results contribute to this debate by demonstrating the presence of two distinct clusters in the superior frontal region, i.e. C7 (putative area 8B) and C8 (putative area 9), separated by the posterior supraprincipal dimple, consistent with cytoarchitectonic maps (Figure 1 E and 3 B). Thus, the intrinsic FC supports the distinction of this region into an area 9 and a distinct area 8B, in line with the Walker (1940) and Petrides and Pandya (1994) parcellation schemes.

The two clusters exhibit very similar connectivity profiles that assign both to the broad prearcuate group of clusters (Figures 4 and 6). However, certain notable differences are apparent. C7 (area 8B) exhibits pronounced connectivity with high-level visual related areas in the dorsal prelunate gyrus and area Opt at the junction of the parietal with the occipital region, and the cortex in the caudalmost part of the superior parietal lobule close to the parieto-occipital sulcus. In the temporal lobe, the connectivity is primarily involving the temporal visual related region. These findings are consistent with some of the available information about the connectivity of area 8B (case 2 in Petrides and Pandya, 1999; Markov et al., 2014). A mild involvement of auditory-related areas in the pattern of connectivity was also observed which, despite possible contamination due to spatial adjacency with the dorsal part of the inferior temporal cortex, is consistent with invasive tract-tracing findings (Romanski et al., 1999). The aforementioned connectivity pattern was absent from cluster C8 (area 9). Thus, the connectivity profile of C7 (area 8B) suggests a role in visuo-auditory and motor functions, in line with recent electrophysiological findings (Lucchetti et al.,2008).

### The border between dorsal and ventral motor/premotor areas

A classic and widely accepted two-way division of the premotor areas is the dorsal/ventral division (e.g., Matelli et al., 1985; Matelli and Luppino, 2001; Barbas and Pandya, 1987; Hoshi and Tanji, 2007). Such a division is also supported by a theory postulating a dual origin of the neocortex (Sanides, 1970; Yeterian and Pandya, 1990). The imaginary caudal extension of the spur of the arcuate sulcus is considered to be the border between the dorsal and ventral premotor areas (Sanides, 1970; Matelli et al., 1985) (Figure 1 D). However, on a cytoand myeloarchitectonic basis, a distinct cortical area is discernible at the cortex caudal to the spur of the arcuate sulcus. This region is characterized by very large neurons in layer V and has been referred to as area 4C by Barbas and Pandya (1987) (Figure 1 B) in accordance with the parcellation of Vogt and Vogt (1919) (Figure 1 A). This is in contrast to the assignment of this region to the ventro-caudal premotor cortex, also known as area F4 (Matelli et al., 1985). The present results indicate that, on a connectional basis, a distinct cluster, i.e. C3 (putative area 4C), indeed occupies the cortical region below and posterior to the spur of the arcuate sulcus towards the central sulcus (Figure 3 B). Importantly, its whole brain FC classifies it with the dorsal motor/premotor group. It is noteworthy that the border between the face and arm representation in this lateral region is postulated to mark the border between the ventral and dorsal premotor cortex (Sanides, 1970). The border between the face and arm representation in the schema from Matelli and Luppino (2001) is ventral to the spur of the arcuate and nicely corresponds to the border defined by the imaginary caudal part of the principal sulcus (Figure 1 D). In conclusion, cluster 3 (putative area 4C) appears to be distinct from the ventral motor/premotor region (Figures 4, 5 and 6). Consequently, our results place the border between the dorsal and ventral motor/premotor areas in the imaginary caudal extension of the principal sulcus. Further evidence from invasive gold standard methods are needed to establish with more certainty the border between the dorsal and ventral premotor cortex, for instance by performing the hierarchical clustering currently employed but with connectional data from invasive tract-tracing cases involving the whole dorsoventral extend of the premotor cortex.

### A connectivity-defined cluster in the fundus of the ventral compartment of the inferior arcuate sulcus as putative area 44

The traditional macroscopic division of the arcuate sulcus is into a superior arcuate sulcus and an inferior arcuate sulcus (Paxinos et al., 2008). Recent examination of the inferior arcuate sulcus in many brains has demonstrated a consistent sigmoid-like shape, which often divides into a clear dorsal and a ventral compartment (cases depicted in Figure 2 in Petrides and Pandya, 2009; cases in Frey et al., 2014). A macroscopic division of the inferior arcuate sulcus into a dorsal and a ventral part is also discernible in the F99 template that we have used in the present study (dias and vias in Figure 2 A). Our connectivity-based parcellation demonstrates the presence of two distinct clusters occupying the cortex within the ventral (vias) and dorsal (dias) parts of the inferior arcuate sulcus, respectively. These are clusters C14 (putative area 44) and C19 (putative FEF region) (Figures 3 B, 7, 9) (see also below) and are shown to be clearly distinct from prearcuate clusters C12 and C13 and postarcuate clusters C15, C17, and C18 (Figure 3 B).

In the human brain, immediately anterior to the ventral part of the premotor cortex (area 6), which is involved with the control of the orofacial musculature, lies a distinct area known as 44 which has been shown to be a critical component of the region involved in language production (Broca’s region). Earlier attempts to identify a homologue of area 44 had considered that the macaque area F5, which is found on the postarcuate cortical region just caudal to the inferior arcuate sulcus, and its anterior extension (F5a) into the posterior bank of the inferior arcuate sulcus may be a homologue of area 44 in the human brain (Geyer et al., 2012). These suggestions were largely driven by an attempt to relate the classical mirror neuron findings in premotor area F5 to the development of language. On the basis of a cytoarchitectonic comparison of human and macaque monkey cortex, Petrides and Pandya (1994) considered a region immediately anterior to premotor area F5 (also referred to as 6VR) in the depth of the ventral part of the inferior arcuate sulcus as a homologue of area 44 of the human brain. Later, this region was shown to be involved with the orofacial musculature based on microstimulation and single neuron recording (Petrides et al., 2005). The connectivity of this region in the macaque monkey was recently clarified by invasive anterograde and retrograde tract tracing studies (Petrides and Pandya, 2009; Frey et al., 2014).

The results of the present study clearly demonstrate the presence of a distinct cluster in the fundus of the ventral part of the inferior arcuate sulcus, namely C14 (Figures 3 B and 7) which is clearly differentiated from the adjacent anterior prearcuate cluster (C12 possibly corresponding to area 45) and two distinct clusters on the posteriorly adjacent ventral premotor cortex, namely clusters C15 and C17, which appear to correspond to two distinct post-arcuate areas previously referred to as area putative ProM (6bβ in the terminiology of Vogt and Vogt, 1919) and a distinct part of he ventral part of premotor area F5/6VR that appears to correspond with area 6bα of Vogt and Vogt (1919), respectively. Hence, our results corroborate previous histological findings by demonstrating, on a connectional basis, the presence of a distinct cluster within the ventral part of the inferior arcuate sulcus colocalizing well with what has been previously described as area 44.

The FC of C18 in the ventral part of the inferior arcuate sulcus corresponds largely with that examined with anterograde and retrograde methods in the macaque monkey (Petrides and Pandya, 2009; Frey et al., 2014). The discovery of area 44 in the depth of the ventral part of the inferior arcuate sulcus of the macaque monkey has generated a debate as to the pre-linguistic role of this area and its recruitment for the control of certain aspects of language production with the evolution of language in the human brain (Petrides, 2006). Recently, Conde et al (2011) demonstrated neurons in the ventral part of the premotor cortex that are involved in the voluntary control of phonation, an important component in the neural machinery necessary for the emergence of language. It has been argued that area 44 is a specialized area that lies between the ventral premotor region that controls orofacial and manual action and prefrontal areas involved in the controlled retrieval of information from memory and may thus have been in a privileged position to mediate between information retrieval and communicative action (Petrides, 2006). Retrieval of information necessary to respond to a specific need would be necessary before action could be organized to convey the subject’s communicative response. Thus, a pre-linguistic neural circuit centered around area 44 might have been ideally suited to the needs of language expression as language evolved.

Immediately posterior to C14 within the ventral part of the inferior arcuate sulcus, we identified two clusters C15 and C17. C15 appears to correspond well with 6bβ of Vogt and Vogt (1919) and 6Vb of Barbas and Pandya (1987), a region of the cortex that has also been referred to as area ProM. C17 appears to correspond with 6bα of Vogt and Vogt (1919) and 6Va of Barbas and Pandya (1987) and electrophysiological data in the macaque monkey has suggested that this ventral oblique strip of cortex may represent the laryngeal/vocal musculature part of the cortex (Hast et al., 1974; Simonyan and Jürgens, 2005).

In conclusion, our present findings, in conjuction with earlier anatomical and physiological research in the macaque monkey suggest the existence of an orofacial dominated region in the ventral part of the inferior arcuate sulcus (C14/area 44) that is surrounded posteriorly by a strip of cortex that represents the orofacial/vocal musculature (C17/area 6bα or 6Va) and posteroventrally by another strip C15 (area ProM or 6bβ or 6Vb).

### A connectivity-defined cluster in the fundus of the dorsal compartment of the inferior arcuate sulcus as putative area 8Av/FEF

In the dorsal part of the inferior arcuate sulcus another cluster was identified based on resting state FC, namely C19 (putative FEF). This area appears connected with visual-related areas, in sharp contrast to the connectivity of C14 (putative area 44) that exhibits a somatomotor profile (Figure 8). It is noteworthy that this sharp connectivity distinction is reflected in the effects of intracortical microstimulation:neurons in the fundus of the ventral ramus of the inferior arcuate sulcus elicit orofacial responses whereas neurons in the cortex in the dorsal ramus of inferior arcuate sulcus, mostly located in its rostral bank, elicit occulomotor responses (Petrides et al., 2005) (Figure 9). Although traditional accounts often link the frontal eye field region with granular prefrontal area 8Av, there is strong evidence that on a microstimulation basis this region lies in the fundus and anterior bank of the arcuate region, a region where a transition between agranular area 6 and the fully granular cortex of the prearcuate cortex takes place. Furthermore, functional neuroimaging evidence in the monkey (Koyama et al., 2004; Savaki et al., 2014) indicate that the premotor agranular cortex in the caudal bank of the arcuate sulcus is also involved in occulomotor function. Such topological characteristics of the FEF are consistent with the location of C19.

**Figure 9.**
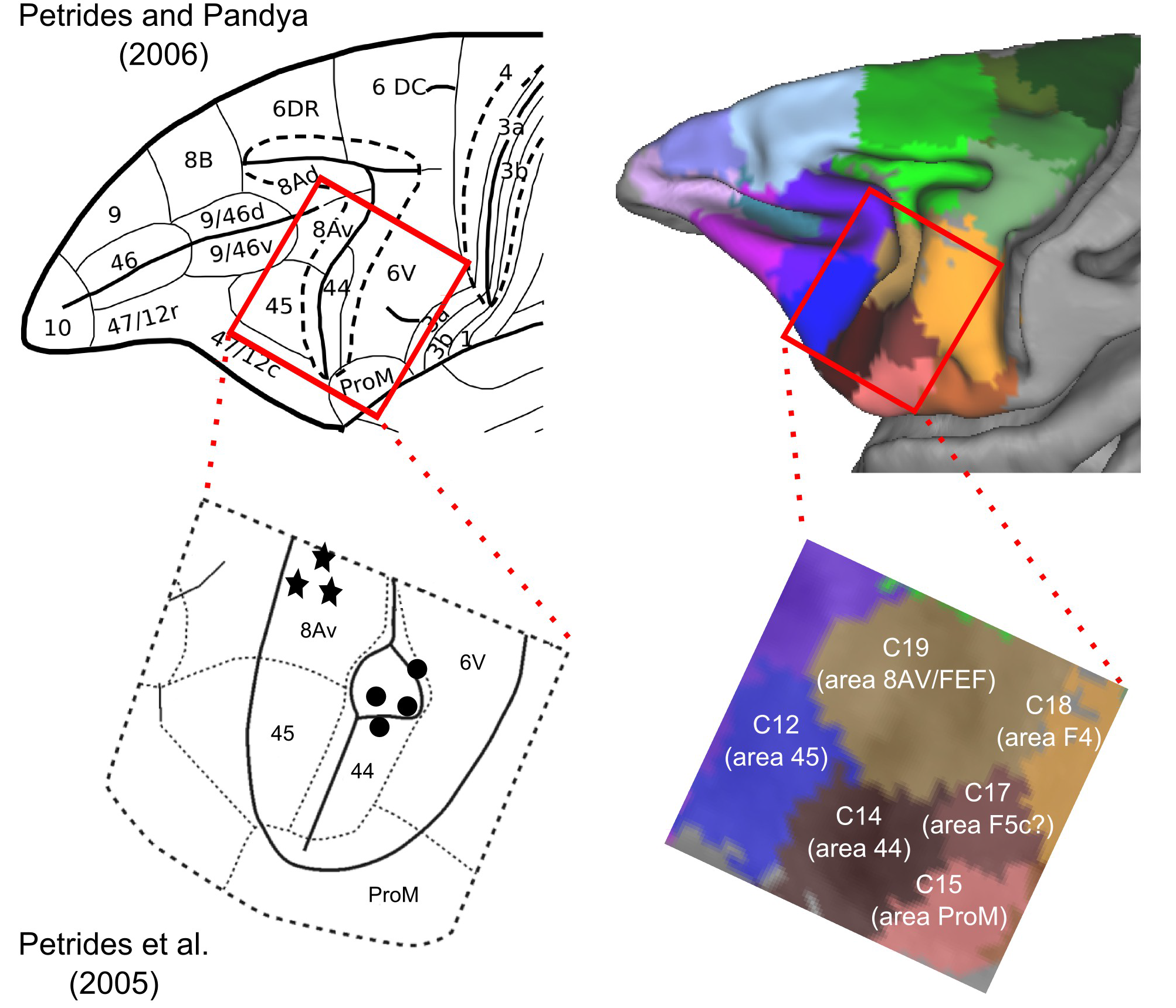
Cytoarchitectonic maps of motor and premotor cortical areas. The red box represents the zoomed-in images of the inferior arcuate sulcus region at the bottom. The circles and stars represent approximate locations of orofacial and occulomotor neurons as identified by intracortical stimulation (Petrides et al., 2005). Note that the occulomotor and orofacial neurons correspond to distinct connectivity-based clusters identified in putative areas 8Av/FEF and area 44 respectively.

In conclusion, the parcellation results of the present study contribute to a resolution of ambiguities concerning the ventral extent of the occulomotor related cortex within the inferior arcuate sulcus, indicating the presence of two areas with distinct connectional profiles that occupy distinct macroscopic subdivisions of the inferior arcuate sulcus. Thus, the macroscopic distinction, vias and dias, might be used for an approximation of the borders of these two areas. Moreover, the current connectivity-based map, within the limitations of rsfRMI, offers putative borders of these two areas with the adjacent post– and prearcuate regions, as well as their whole brain connectivity similarity (Figures 8 and 9).

### Limitations and perspectives

Connectivity, estimated from rsfMRI data, does not provide the level of detail of gold standard invasive tract-tracing techniques. In addition, connectivity maps estimated from rsfMRI might include areas between which no direct anatomical connectivity exists, reflecting polysynaptic connectivity (Adachi et al., 2012; Goñi et al., 2014). A much needed next step is to assess quantitatively the degree of correspondence of FC and connectivity estimated with invasive tract-tracing methods (see Miranda-Dominguez et al., 2014 for a first attempt of such quantification). Despite the limitations in specificity and resolution of the method, the present results inform current debates about the organization of the LFC by providing evidence of connectional divisions of the LFC. The precise boundaries, on a cytoarchitectonic basis, of these divisions and their precise connectivity pattern can be uncovered by quantitative cytoarchitectonic analysis (e.g. Mackey and Petrides, 2010).

The clusters currently uncovered could consist of further subdivisions. For instance, C5/area F7/6DR is usually treated as non-homogenous, consisting of a supplementary eye field and a non-supplementary eye field zone (Luppino et al., 2005). The organization of the cortex could be represented as a hierarchy, spanning several topological scales (Meunier et al., 2010). Indeeed, broader divisions of the LFC have been found using the rsfMRI data (Hutchison and Everling, 2013). Higher spatial resolution alongside with advancements in clustering approaches, despite substantial challenges (see Lancichinetti and Fortunato, 2011), could potentially offer a more fine grained parcellation, moving towards lower spatial levels of the LFC organization.

Signal contamination between adjacent banks of sulci, such as the dorsal and ventral bank of the principal sulcus, render problematic the accurate parcellation of cortical areas within sulci. Such limitations might be circumvented with higher spatial resolution during the rsfMRI acquisition. The dorsal and ventral LFC clusters that are the focus of the current study are largely not influenced by such signal contamination with C19 being an exception since it might also encompass parts of the ventral premotor areas in the posterior bank of the arcuate sulcus.

### Conclusions

We investigated the intrinsic functional architecture of the LFC of the macaque in order to elucidate current debates on its architecture. Within the dorsal LFC, we demonstrate that i) the posterior supraprincipal dimple constitutes the border between two areas ii) area 6DC/F2 contains two distinct connectivity-defined areas and iii) a distinct area exists around the spur of the arcuate at the border of the dorsal/ventral division of the LFC. Our results within the ventral LFC clearly demonstrate the presence of a putative area 44, with a somatomotor/orofacial connectional signature. This area is located in the fundus of the vias and is differentiated from premotor and prearcuate clusters, bounded dorsally by a distinct cluster with an occulomotor connectional signature, identified as putative FEF and located in the dias. Both of these areas were discernible from premotor clusters. The current map can be used for future cross-species examination of putative homologues in the human LFC with the aid of the same modality, namely rsfMRI.

## Acknowledgements

We would like to thank Laura Wallor for assistance in preparing Figure 1.

